# Robust scan synchronized force-fluorescence imaging

**DOI:** 10.1101/2020.05.15.098970

**Authors:** Patrick Schmidt, John Lajoie, Sanjeevi Sivasankar

## Abstract

Simultaneous atomic force microscope (AFM) and sample scanning confocal fluorescence microscope measurements are widely used to obtain mechanistic and structural insights into protein dynamics in live cells. However, the absence of a robust technique to synchronously scan both AFM and confocal microscope piezo stages makes it difficult to visualize force-induced changes in fluorescent protein distribution in cells. To address this challenge, we have built an integrated AFM-confocal fluorescence microscope platform that implements a synchronous scanning method which eliminates image artifacts from piezo motion ramping, produces intuitive, accurate pixel binning and enables the collection of a scanned image of a sample while applying force to the sample. As proof of principle, we use this instrument to monitor the redistribution of fluorescent E-cadherin, an essential transmembrane protein, in live cells, upon application of mechanical force.

## Introduction

Integrated force-fluorescence microscopes, which consist of an atomic force microscope (AFM) mounted on an inverted epifluorescence microscope, are widely used to image the topography of cells while simultaneously visualizing the distribution of fluorescently tagged proteins within the cell (*1-5*). While these microscopes are useful for locating a point of interest (with fluorescence imaging) and then probing that point of interest (with AFM), they cannot be used to monitor a force-induced fluorescence response, such as the redistribution of cellular proteins. This is because the large field of view in a widefield fluorescence microscope images (and potentially photobleaches) many cells, while only a single point on a single cell surface is simultaneously probed with the AFM. Furthermore, due to limited optical sectioning capabilities of many widefield setups, background information away from the focal plane makes it impossible to determine where fluorophores reside along the optical axis, and consequently cannot be used to image a section of interest, such as the cell’s surface. While total internal reflection fluorescence microscopy can be used to collect images from a thin optical slice, the fluorescence collected is typically limited to the region near the cell-substrate interface and as such the apical surface of a cell cannot be visualized.

Point scanning confocal microscopes are better suited to image thin optical sections in cells with high resolution. In these microscopes, images are acquired either by scanning the sample through a fixed laser position or scanning the source laser through an area with the use of galvo mirrors while keeping the sample fixed. The latter has the benefit of leaving the sample stationary and opens the possibility of simultaneously probing the sample during image acquisition. However, moving the beam relative to the input of a powerful objective causes optical aberration and the inherent sinusoidal motion of the galvo mirrors causes image distortion. A stationary sample also cannot be stabilized with active feedback, which is useful for monitoring a point of interest over a long period of time and reducing unwanted sample drift (*6, 7*). In contrast, sample scanning confocal microscopy (SSCM) is ideal for fluorescence imaging of the apical cell surface. However, in an integrated SSCM-AFM instrument, the AFM tip cannot be used to apply force to a predefined point on a sample unless the AFM tip’s lateral motion is synchronized with sample scanning.

To accomplish synchronized scanning of an AFM and SSCM, we recently developed a custom AFM which utilizes a piezo stage capable of long range (100µm) motion in all three spatial axes (*8*). Here, we integrate this AFM with a SSCM and present a method to acquire images while simultaneously applying a force with the AFM tip at a predefined point, without any piezo ramp aberrations. We demonstrate the capabilities of our method by directly imaging the effect of force on the apical cell surface distribution of a fluorescently tagged E-cadherin, an essential transmembrane adhesive protein (*9, 10*).

## Methods

Our instrument (Figure 1) utilizes a typical sample scanning confocal fluorescence setup (*11*) with a three-axis piezo sample stage (P-733.3, Physik Instrumente) mounted on a modular microscope frame (RAMM, Applied Scientific Instrumentation). A 532nm laser (OBIS, Coherent) couples into a single mode fiber (SMF). The beam is collimated at the fiber endpoint (C) and reflected by a mirror (M) towards the 60x microscope objective (O, Olympus) and focused at the sample mounted on an XYZ piezo stage. The fluorescence and a fraction of the backscattered light are collected back through the objective and diverted by a dichroic (D1, Chroma). A second dichroic (D2, Chroma) splits the backscattered and fluorescent light which are collected by avalanche photodiodes (APD 1 & 2, respectively, Micro Photon Devices). A field programmable gate array (FPGA, National Instruments) is used to command piezo stage position, and acts as a photon counter for the APDs and partially processes the data (with the rest done on the host PC) to form the resulting scan images.

**Figure 1:**
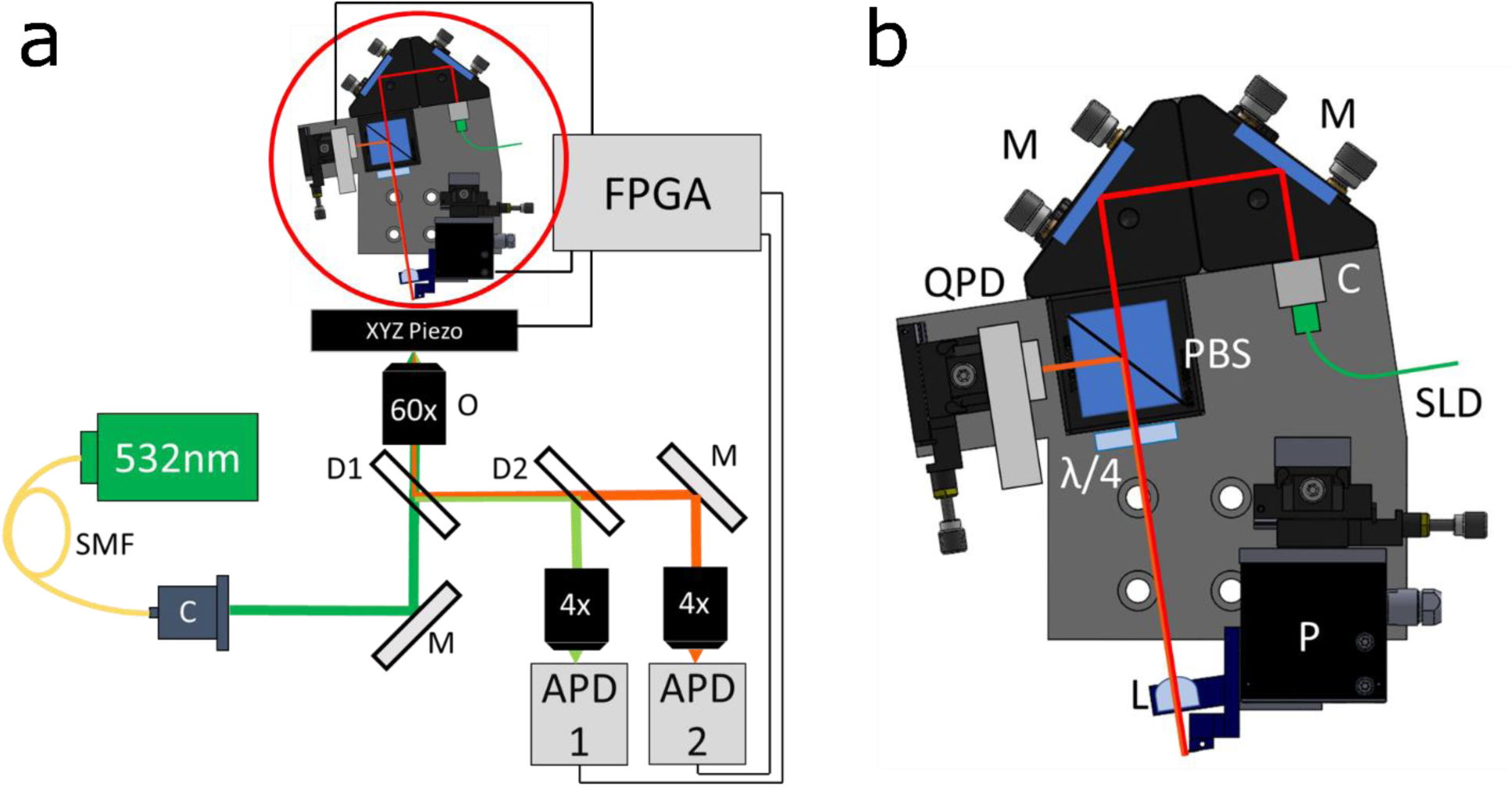
Instrumentation. **(a)** Diagram of microscope, SMF- Single mode fiber, O- objective, M- mirror, D1/D2- dichroic, C- collimator, FPGA- field programmable gate array, APD1/APD2- avalanche photodiode. **(b)** Diagram of custom AFM. M- mirror, L- lens, C- collimator, SLD- superluminescent diode, QPD- quadrant photodiode, PBS- polarizing cube beamsplitter, λ/4- quarter waveplate, P- 3 axis piezo.

The AFM is situated above the microscope sample (circled in red) and shown in detail in Figure 1(b). A fiber-coupled infrared superluminescent diode (SLD, 850nm, QPhotonics) mounts to a collimator (C, Thorlabs) which in turn mounts to two adjustable right-angle mirrors (M, Thorlabs), allowing for fine positioning and angling of the beam onto the AFM cantilever. The beam passes through a polarizing cube beamsplitter (PCB, Thorlabs) and a quarter waveplate (λ/4, Thorlabs) resulting in a circularly polarized collimated beam. This is focused through a lens (L, Newport) on to the AFM cantilever. Upon reflection, the polarization handedness flips, resulting in the reflected light being polarized perpendicular to the incoming beam, and as such is reflected by the PCB to the quadrant photodiode (QPD, First Sensor). The AFM chip (ARROW TL2, NanoWorld) is mounted at an 8.5° angle on the three-axis piezo stage (P-616, Physik Instrumente) to allow it to probe a surface, and the rest of the optics are mounted to match that angle for perpendicular reflection. The QPD signal is digitized by on board analog to digital converters (ADCs) on the FPGA, which in turn can control the motion of the piezo stage. Each piezo stage is controlled with PI’s E-712.3 controller.

To test our synchronized force-fluorescence microscope we designed a cell pressing assay. We wanted to monitor change in the distribution of cell membrane proteins due to force-induced deformation of a cell, so we used Madin-Darby Canine Kidney (MDCK) cells expressing DSRed-tagged E-cadherin (a surface protein integral in mediating cell-cell adhesion). We grew these cells on a piranha (3:1 H2SO4:H2O2) -cleaned glass coverslip which was mounted to our fluorescence microscope piezo. In order to apply force on these cells without puncturing their membrane, we positioned borosilicate microspheres (diameter < 10µm, Polysciences) at the ends of tipless AFM cantilevers (Arrow TL2, NanoWorld) using a micromanipulator (CellTram, Eppendorf). These microspheres were then sintered to the cantilever (*12*) in an oven (Isotemp, Fisher Scientific) using a slow heat ramp (room temperature to 500°C at 8°/min, 500°C to 800°C at 5°/min, hold at 800°C for 1 hour). Since the microspheres are of a scale comparable to the size of MDCK cells, they do not puncture the cells even with a large applied force. The technique used to synchronously scan the AFM and confocal microscope piezo stage and the results of the cell force-fluorescence application are detailed in the results section.

## Results

The AFM and fluorescence microscope piezo stages cover a 2D space by scanning the piezo along one axis and stepping it across the second axis in between scanned lines (Figure 2). Smooth motion along the scan axis was achieved by commanding the piezo to move in a positional “ramp”, that is move through the scan line with a constant velocity. To save scanning time and minimize undesired motion, the direction of the ramp was switched with each scan line (Figure 2). Each scan line was distributed across the requested area in such a way that each resulting image pixel represents light collected from the center of that pixel (rather than the edge or corner, Figure 2) yielding a more intuitive sense of the scanned space and a better ability to locate the absolute position of features.

**Figure 2:**
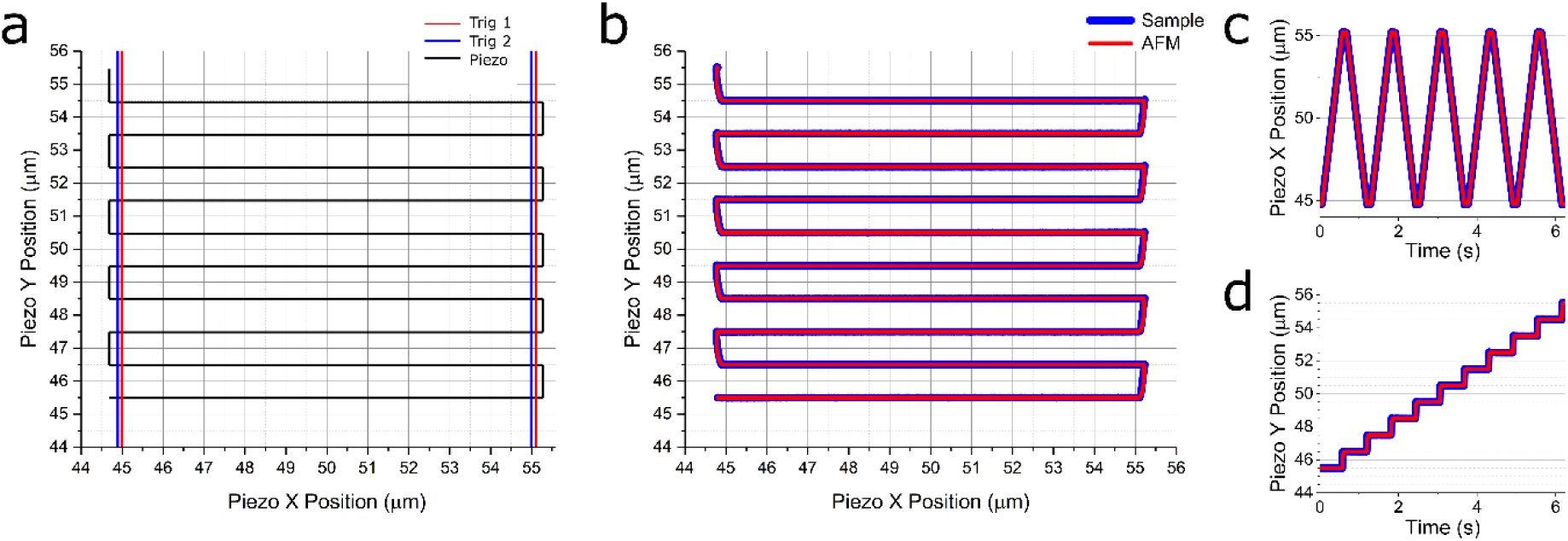
Scanning method and synchronization. **(a)** A 2D lateral scan is defined over a 10×10µm area with 10 scan lines. The scan definition is such that when the resulting data is formatted into a 2D image, each pixel value represents light collected from the center of that pixel. The scanning axis (X in this case) also extends before and after the scan area of interest to allow the triggering system which eliminates “warping” at the edges. Note that the range for each trigger extends slightly beyond the scan area, allowing a trigger “hand off” between scan lines such that the trigger bounce does not signal the start of the next scan line collection. **(b)** Each piezo (sample and AFM) are programmed to internally record their positions and report the result at the end of the scan. To be sure the scan is not only spatially, but also temporally synced, the 2D scan is separated into each axis and plotted against time, X shown in **(c)** and Y shown in **(d)**. The traces verify that the two piezo stages are synchronized both spatially and temporally.

While the scan method described above ideally evenly covered the requested 2D space, there was a ramp up and ramp down at either end of the piezo stage travel which warped the image. This undesired effect limited the temporal resolution since it scaled with the time of each scan line, in that over the same scan length a faster scan appeared more warped. To address this problem, we programmatically adjusted a user defined scan line to include scan “padding,” that is, a short distance before and after the requested scan dimensions was built in, and the scan time was adjusted to maintain the requested velocity over the requested scan distance. The padding was chosen to allow enough time and space for the piezo to reach target velocity before reaching the scan line start, and enough time after to reduce velocity to zero before starting the next scan line. The data from these ramp up/ramp down regions were then faithfully excluded from the resulting image. If the piezo stage has a consistent response time, these regions could be excluded by discarding a set amount of data collected over the ramp up/ramp down time and generating the image using the remaining data. However, since this time varies between piezo stages, and even between different load weights on the same stage, we relied on the recorded piezo position, using digital position triggers to exclude unwanted ramping data. We set a trigger output to be high only over the user defined scan line (excluding the padding), and only collected detector data when the trigger was high. In this configuration it became necessary to address the problem of digital trigger bouncing (that is rapid changing from high to low for a time before stabilizing on its final state), since it breaks image acquisition. Typical “debouncing” solutions eliminate this problem at the cost of time resolution, which would also be detrimental to our images. For this reason we implemented a two trigger scheme, one indicating the start of the scan line and the other indicating the end of the scan line (Fig. 2(a)). This way we used the first switch from each trigger and ignored altogether the bounces that follow.

To faithfully scan the two piezo stages over the same area, synchronized in time and space, the two piezos’ closed-loop motion parameters were tuned such that when the same motion command was sent to both stages, their response time and settling time matched. However, even with their motion tightly coupled, the effort is rendered moot if the scan does not start at the same time for both piezo stages. When the command to start the scan was sent from the host computer (via USB), we saw significant discrepancies in the actual scan start time, resulting in an offset between the two stages for the duration of the scan. To address timing differences, we first synchronized the two controllers’ clocks via cable, ensuring commands received at the same time will execute at the same time. To ensure the commands arrive at the same time, we set the pre-defined scan waveforms to execute when a third digital trigger is set high, sent via FPGA digital output to both piezo controllers. Consequently the controllers received and executed the scan command at the same time.

These features combined allowed an AFM tip to scan synchronously with a feature of interest on the sample to generate a faithful backscatter and fluorescent image. Because this only affects the lateral motion of each stage, the optical Z axis of the AFM is free to move independent of the scan, which will eventually allow force clamp experiments while imaging is taking place. More immediately, this allowed us to image the AFM tip and sample fluorescence simultaneously while maintaining a user-controlled AFM pressing distance which is useful for gaining mechanistic insight into pliable substances such as cells.

To verify the scan synchronization we glued one AFM cantilever to a glass coverslip which was mounted to the sample piezo stage. We then brought a second AFM cantilever down near the coverslip with the custom AFM apparatus. We defined a scan and commanded both the sample and AFM piezos to start moving as described above. We recorded their motion with a camera inserted into the detection arm of the microscope (Supplementary Video). Our data showed that the cantilevers maintain their position relative to one another while scanning through the 2D space, over the laser focus. This shows the scans begin at the same time and maintain registry throughout the scan. To quantify this we also record the piezos’ positions during such a scan, shown in Figure 2(b-d).

With scan synchronization verified, we next used the platform to simultaneously image and probe live cells, prepared as described in the Methods section. Viewing the cells and AFM cantilever through the camera, we initially roughly aligned a group of cells over the focused laser using manual micrometer actuators on the microscope stage. The objective height was adjusted such that the top of the cells were in focus. We then similarly aligned the AFM cantilever over the cells with actuators on the custom AFM. Next an initial large synchronized scan was commanded, and a wide view of the cell fluorescence and AFM cantilever and microsphere were generated. Next, we selected a new scan center points on the fluorescence and backscatter images to redefine the sample and AFM scans, respectively; a cluster of fluorescent E-cadherin was chosen for the sample, and the center of the microsphere was chosen for the AFM. The result of the following 50×50µm scan is shown in Figure 3(b,c). Keeping this scan center, we zoomed in on a fluorescent E-cadherin cluster and microsphere by commanding a smaller scan area. We then commanded the AFM piezo to move the cantilever toward the sample until a small deflection was detected from the AFM QPD, indicating the cantilever had made contact with the cell. We then applied progressively higher force to the cell by moving the AFM piezo toward the sample, then reduced force by moving away, collecting images throughout (Figure 3(e,f)). As the AFM cantilever moved into the focused waist of the confocal laser beam, it backscattered more light, resulting in a brighter image. At the same time, a ‘dark spot’ formed in the fluorescent cell image, which corresponds in shape and position to the AFM microsphere. We interpret this ‘dark spot’ as E-cadherins on the cell surface being pushed out of the laser focus by the microsphere. Indeed, as the microsphere was lifted from the cell, fluorescence recovery was observed at the pressing site. This result demonstrates the use of this synchronized force-fluorescence platforms in simultaneously imaging and manipulating biological samples.

**Figure 3:**
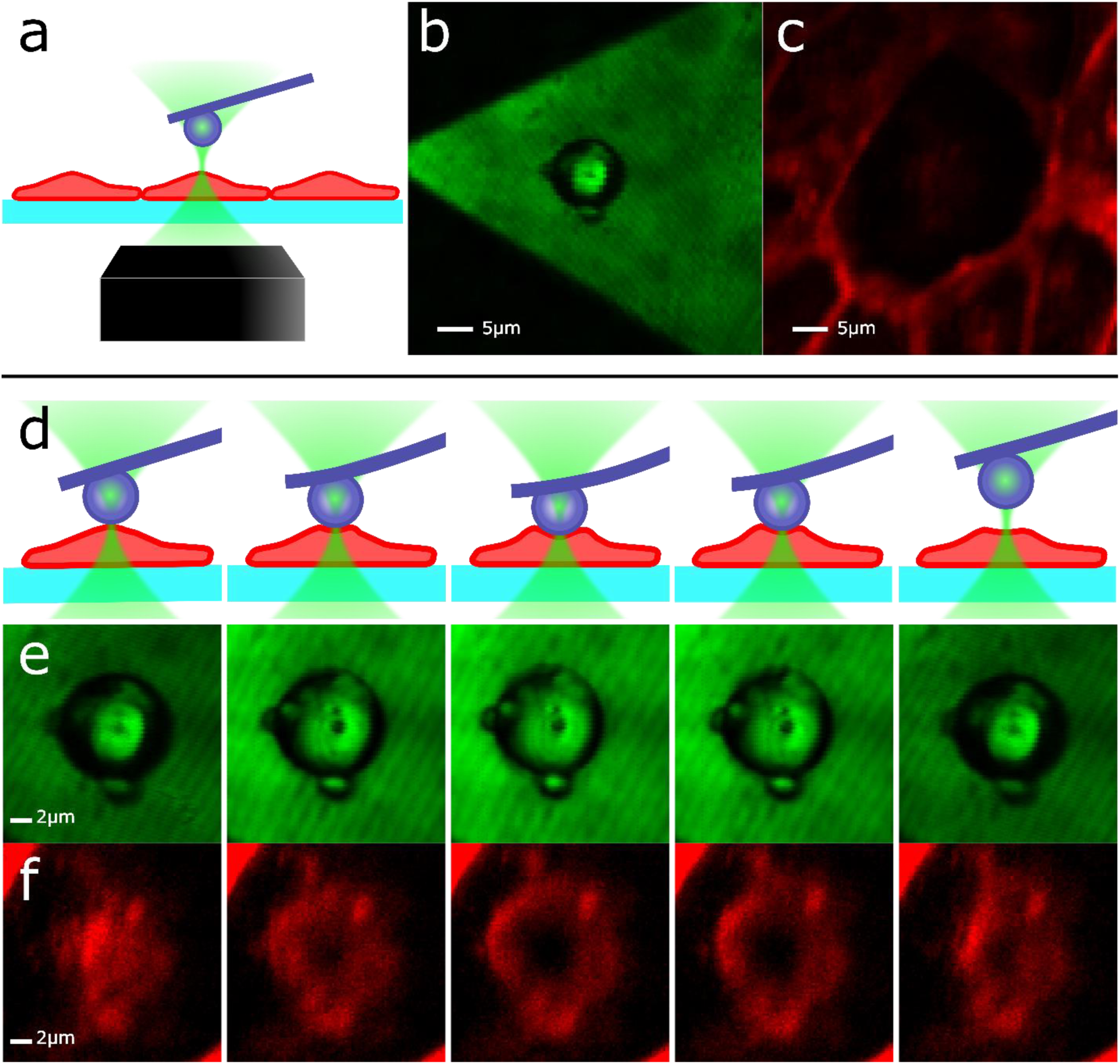
Imaging force-induced redistribution of cell surface proteins. To demonstrate the utility of the synchronized force fluorescence platform, we designed an experiment in which we collect fluorescence and backscatter images while applying a varying force on the apical surface of cells expressing DSRed-tagged E-cadherin. **(a)** Illustration of the experimental setup. The objective focus is set to the apical surface of the cell of interest (cell shown in red, beam waist in green), and the AFM cantilever with microsphere is positioned over the laser focus (or anywhere within the scan area the user chooses). **(b)** Backscatter and **(c)** fluorescence image results of a 50×50µm scan from APD 1 & 2, respectively (Figure 1). The triangular shape of the AFM cantilever as well as the silica microsphere sintered to the cantilever (along with a few smaller microspheres) are visible in *(b)*. The fluorescently tagged E-cadherin largely reside at cell-cell junctions, making the edges of the cells bright as seen in *(c)*. **(d)** Progression of the force applied to the cell by the AFM, and the resulting **(e)** backscatter and **(f)** fluorescent images (20×20µm scans). In row *(e)* we see the microsphere centered in the image, and as it presses into the cell the cantilever becomes brighter as it moves into the laser beam waist. In *(f)* we see a cluster of fluorescent E-cadherin on the apical surface of a cell. As higher force is applied, these cadherin are pushed out of the beam waist, and we see a dark spot form where the microsphere is pressing into the cell. Upon AFM withdrawal, we see the fluorescent E-cadherin return to the pressing site as the cell reforms its original shape. These images serve as good confirmation that our synchronized scanning method allows fluorescence imaging while applying force.

## Discussion

In this paper, we have demonstrated the design and use of our novel synchronized force-fluorescence confocal scanning microscope. By taking care to ensure the area scan is accurate, repeatable, and synchronized between two piezo stages, we can image the force response of living cells through redistribution of fluorescently tagged membrane proteins. We believe this instrument has great promise in a variety of force fluorescence applications, mainly in imaging molecules and cells during force application with higher resolution and greater optical axis range than existing methods.

In these experiments, we apply a very large force (between 1 and 90nN) to effect a clear change in our fluorescence image. This force range may also be useful in determining cell and tissue mechanical properties, such as cell viscoelasticity (*13, 14*) (Supplementary Figure S2). For more delicate experiments, applying lower forces is possible. Our custom AFM has a force resolution of 6pN at an acquisition bandwidth of 1kHz (*8*) for measurements conducted in air (Supplementary Figure S1). However, in an aqueous environment, our AFM force resolution is limited by the reduction of light intensity falling on the QPD (72pN, Supplementary Figure S1). This limitation can be easily overcome by replacing the superluminescent diode with a more powerful light source. Of course if the light source is too bright, a fluorescent sample will photobleach, so that balance will need to be struck with further experimentation.

Another issue arises when monitoring the reported force during a 2D scan. Our custom AFM is designed to ideally send the superluminescent diode beam perpendicular to the AFM cantilever, which when aligned correctly should eliminate any change of the beam’s deflection during the cantilever’s lateral scan motion. However the experimental data shows that this alignment is very sensitive, and a small deviation from a perpendicular reflection can cause a change in force signal when no force is being applied during a lateral scan. This is due to the long lever arm of the beam, which provides an advantage for monitoring small deflections, but a disadvantage during scanning. If the beam is not perpendicular to the cantilever and the cantilever moves relative to the beam, we see a resulting shift on the QPD. This can be addressed in a number of ways, perhaps the easiest being a more precise milling of mounting parts with more robust connections to ensure better angle control. Another option would be to largely replace our AFM optics scheme with a more traditional layout, allowing light source, cantilever, and detector to all mount on the piezo stage and move in unison. This would ideally be quickly swappable with our current AFM mount to allow use of both styles on the same fluorescence microscope.

Finally, our work provides a clear path forward to integrate our custom AFM and our previously published fluorescence-compatible active feedback platform (*6*). The next step is to apply our feedback technique on an AFM tip (or microsphere) mounted to our custom AFM apparatus and extend its feedback capability to three dimensions. With our feedback platform working in conjunction with our custom AFM, we’ll be able to perform single molecule manipulation and force measurement in a repeatable fashion on specifically located molecules, where previously these type of experiments relied on stochastically probing single molecules from within a reservoir of such molecules.

## Acknowledgements

We would like to thank Omer Shafraz for assistance with design of the biological assay. Research reported in this publication was supported in part by the National Institute of General Medical Sciences of the National Institutes of Health (R01GM121885).

