## Supplemental data for "Robust scan synchronized force-fluorescence imaging"

### Video

This video visually demonstrates the synchronized motion of the sample and AFM. On the left is the AFM-mounted cantilever, and on the right is an identical cantilever glued to the sample surface. The sample and AFM are roughly positioned near each other and above the focused laser.

**Figure S1**

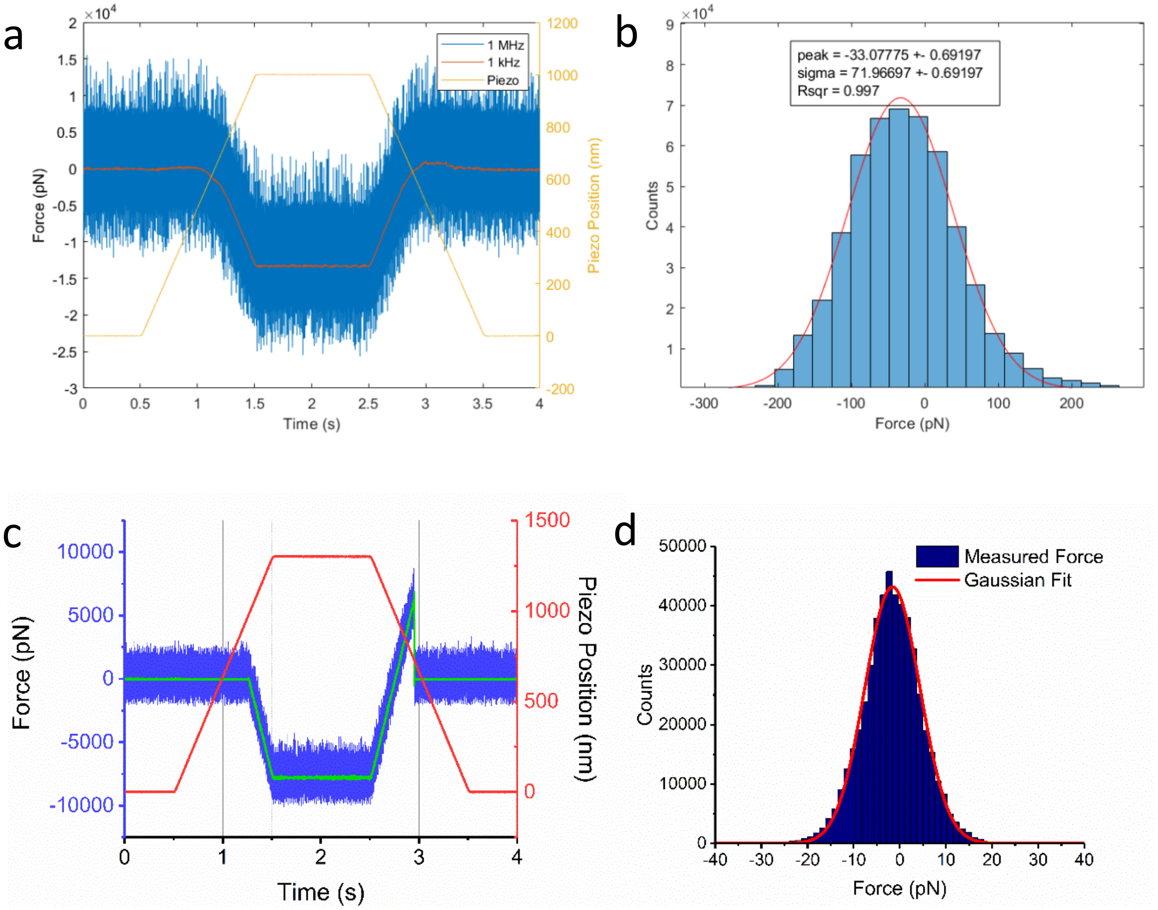


**Custom AFM performance when pressing a glass substrate in water and air. (a)** Full bandwidth force measurement in water (blue) and the same data smoothed to 1kHz overlaid (red). The piezo's z position trace is in yellow (right axis). **(b)** Due to the greatly reduced superluminescent diode intensity in water, the signal to noise becomes low, and our sigma at 1kHz is 10x worse (72pN) than our performance in air. The signal to noise ratio can be easily increased using a brighter superluminescent diode. **(c)** Full bandwidth force measurement in air (blue) and the same data smoothed to 1kHz overlaid (green). The piezo's z position trace is in red (right axis). **(d)** The sigma at 1kHz is 7pN (*data in c and d were previously reported in a conference proceeding: P. Schmidt, B. Reichert, J. Lajoie, S. Sivasankar, Adaptive atomic force microscope. SPIE BiOS (SPIE, 2020), vol. 11246*.).

**Figure S2**

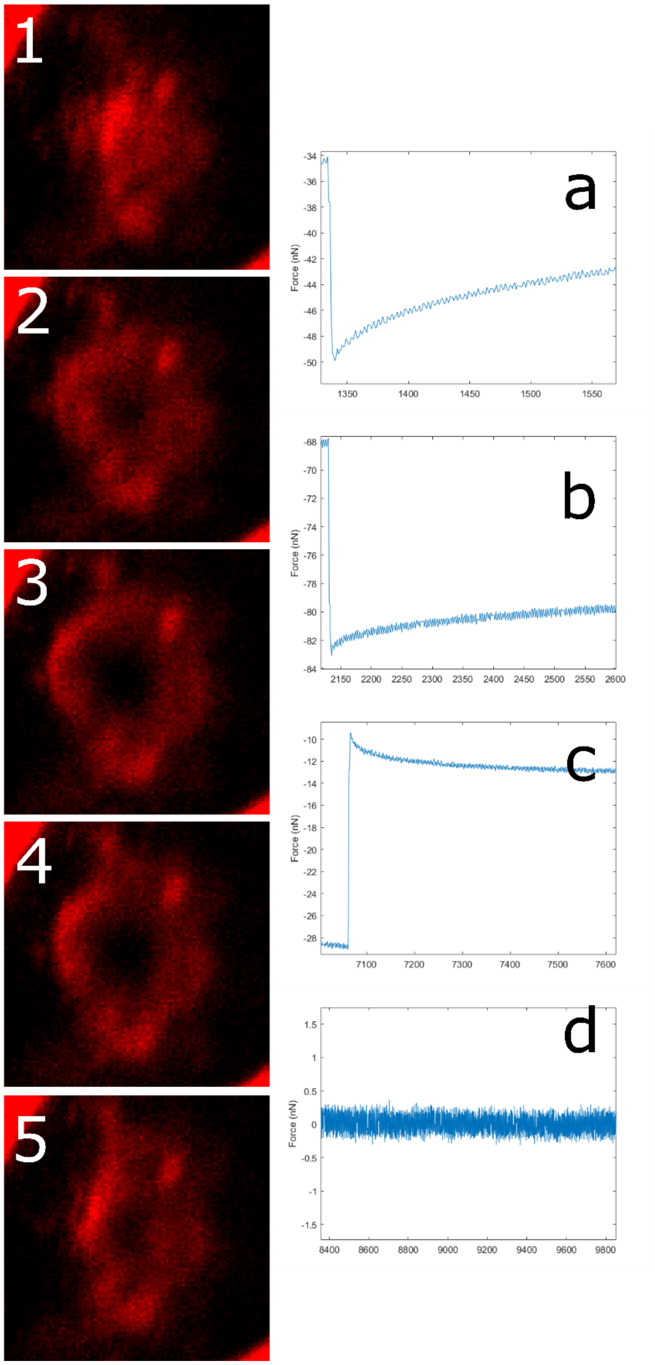


**Simultaneous force and fluorescence measurements.** Column 1 shows the fluorescent images collected for Figure 3, labeled numerically in sequential order. Column 2 shows the corresponding force trace before images 2-5 were acquired. **(a)** shows the AFM force trace before image 2 was acquired, **(b)** before image 3, **(c)** before image 4, and **(d)** before image 5 (flat because the AFM microsphere was no longer in contact with the cell). These force-curves show a characteristic relaxation after piezo motion stops, which can be used to determine the cell's viscoelastic properties.
